# Effect of X-ray irradiation on ancient DNA in sub-fossil bones – Guidelines for safe X-ray imaging

**DOI:** 10.1101/057786

**Authors:** Alexander Immel, Adeline Le Cabec, Marion Bonazzi, Alexander Herbig, Heiko Temming, Verena J. Schuenemann, Kirsten I. Bos, Frauke Langbein, Katerina Harvati, Anne Bridault, Gilbert Pion, Marie-Anne Julien, Oleksandra Krotova, Nicholas J. Conard, Susanne C. Muenzel, Dorothée G. Drucker, Bence Viola, Jean-Jacques Hublin, Paul Tafforeau, Paul Tafforeau

## Abstract

Sub-fossilised remains may still contain highly degraded ancient DNA (aDNA) useful for palaeogenetic investigations. Whether X-ray computed [micro-] tomography ([μ]CT) imaging of these fossils may further damage aDNA remains debated. Although the effect of X-ray on DNA in living organisms is well documented, its impact on aDNA molecules is unexplored.

Here we investigate the effects of synchrotron X-ray irradiation on aDNA from Pleistocene bones. A clear correlation appears between decreasing aDNA quantities and accumulating X-ray dose-levels above 2000 Gray (Gy). We further find that strong X-ray irradiation reduces the amount of nucleotide misincorporations at the aDNA molecule ends. No representative effect can be detected for doses below 200 Gy. Dosimetry shows that conventional μCT usually does not reach the risky dose level, while classical synchrotron imaging can degrade aDNA significantly. Optimised synchrotron protocols and simple rules introduced here are sufficient to ensure that fossils can be scanned without impairing future aDNA studies.

## Introduction

Since its discovery, X-ray imaging has found a broad range of applications in medical, anthropological and palaeontological studies. Conventional X-ray computed [micro-] tomography ([μ]CT) scans are routinely used to generate three-dimensional (3D) models of fossil remains, to explore internal structures, as well as to provide virtual replicas of the fossils that can be shared for analysis with other institutions. Furthermore, CT scanning of fossil remains prior to analysis requiring invasive/destructive sampling has been recommended, in order to preserve valuable internal morphological information^1^. Until now, about half of the sub-fossil remains used for aDNA analysis were scanned before sampling (Supplementary Table 1). However, recent concerns have been raised regarding the potential deleterious effect of X-rays on the retrieval of aDNA^2^, especially in the case of μCT performed using synchrotron sources. These concerns arose originally after the observation of transitory darkening of translucent or white enamel when submitted to high resolution synchrotron scans (typically sub-μm voxel size), that are used to investigate dental microstructures^2,3^. This darkening can be easily removed by using low power, low energy UV (375 nm, ideally from LED), as already demonstrated on hundreds of teeth imaged at the ESRF. Nevertheless it indeed indicates a significant effect of the X-rays on the specimen, mostly due to electronic excitation, but also partially to ionisation^4,5^. One of the most relevant applications of synchrotron μCT, but also the most controversial in regards to the risks for aDNA, is for non-destructive investigations of dental structures and microstructures in palaeoanthropology, using virtual palaeohistology approaches^3,6–8^.

Numerous studies describe the effects of X-ray irradiation on DNA molecules in biological conditions, particularly in medical contexts. Mutations, DNA strand breaks, chemical modifications of bases, and structural changes such as gene order rearrangements have been identified^4,9–11^. Creation and diffusion of radicals during X-ray exposure have been linked to molecular structural modifications^12,13^, since X-ray photons are known to contribute to strand breaks^14^. In living organisms, repair mechanisms exist to withstand moderate levels of radiation-induced damage^15^. In the absence of repair mechanisms, the damages would accumulate with increasing X-ray dose (each new scan adding to the effects of the previous ones). When compounded with additional DNA modifications such as oxidative and hydrolytic damage as a result of natural decay and taphonomic processes^16,17^, downstream genetic analyses may be affected in cases of significant levels of X-ray dose accumulation. Particularly the hydrolytic conditions seem to play a critical role: as demonstrated in a recent study on simulated effects of X-radiation on fragmented DNA in dry, wet and frozen states, the highest probability of radiation-induced DNA damage occurs in a wet state^18^.

Earlier studies have used modern pig (*Sus scrofa*) bones as proxies for archaeological samples, where molecular damage was determined based on the success of Polymerase Chain Reaction (PCR) amplification^19,20^. Preserved bird skins from museum specimens were evaluated via capillary electrophoresis-based DNA quantification to infer the level of fragmentation induced by X-ray exposure^21^. The results of these investigations, however, were not consistent, potentially owing to inappropriate or non-homogenous samples, the use of low sensitivity measures to evaluate DNA concentrations, or insufficient control of X-ray dose. To date, no single investigation has reliably assessed the effects of X-ray irradiation on authentic aDNA molecules. A robust investigation of the effects of accumulation of X-ray dose on ancient DNA integrity was, therefore, urgently needed.

In this study we evaluate the effects of X-ray radiation on aDNA from Late Pleistocene megafaunal teeth and bones. In order to assess the effects of a large range of X-ray dose deposition, we used powerful polychromatic X-rays instead of a conventional X-ray source. Homogenized aliquots of crushed bone and dental tissues were exposed to increasing synchrotron X-ray doses using different exposure times at the European Synchrotron Radiation Facility (ESRF). The dose rate and integrated dose were quantified as water equivalent surface dose for each experimental setup. After exposure, we used both quantitative PCR and next generation sequencing to evaluate the effects of synchrotron irradiation on: (1) aDNA quantity, (2) aDNA fragment length and (3) aDNA-characteristic nucleotide misincorporation patterns.

Our results reliably demonstrate a clear relationship between increasing X-ray dose deposition and level of aDNA damage, likely through increased strand breakage. We observe that strong X-ray exposure can significantly degrade aDNA (surface dose above 10 000 Gray (10 kGy)); however, no effect can be demonstrated for a dose below 200 Gy. Dosimetry on classical synchrotron configurations, as well as on new configurations optimised for low dose imaging used at the ESRF on the beamline ID19 for palaeoanthropology, and finally on two conventional μCT scanners (a BIR ACTIS 225/300 and a Skyscan 1173), demonstrate that in the vast majority of cases conventional microtomographs are well below the detection limit of any defect on aDNA.

## Results

Crushed material from 38 Late Pleistocene animal bones and teeth consisting of bison, cave bear, and giant deer from the Swabian Jura, and roe deer from the French Jura were exposed to various configurations of polychromatic synchrotron beam on the beamline BM05 at the ESRF (see Material and Methods: Table 2). In the first experiment, aliquots were exposed to an extremely high level of X-ray radiation. For the second experiment, aliquots were irradiated with increasing exposure time. In the third experiment, we exposed aliquots to irradiation applicable in conventional high quality imaging μCT.

### Effect of extreme irradiation on aDNA quantity

In the first experiment, aliquots from 11 well preserved Late Pleistocene bones were exposed for 34 minutes to an extreme dose of X-ray radiation of 170 kGy (water equivalent surface dose). Subsequent processing of each sample was performed alongside a non-scanned control, a negative extraction control, and a negative library-preparation control. The amount of aDNA that could be quantified via qPCR decreased substantially in the scanned aliquots compared to the non-scanned controls, indicating a highly damaging effect of this high X-ray radiation dose (Fig. 1). No sequencing data were produced for these exposed samples, as the amplifiable aDNA quantities after scanning were too low, being comparable to the negative controls.

**Figure 1.**
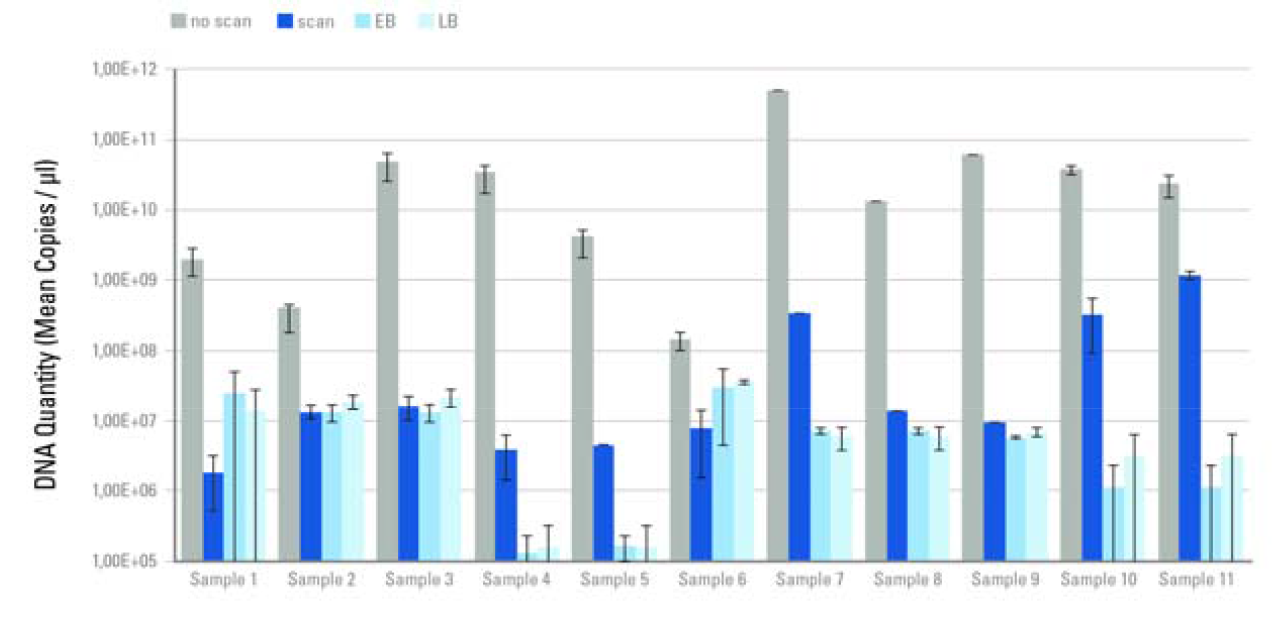
aDNA quantitation of 11 ancient samples after exposure to a radiation dose of 170 kGy. Each column represents a mean copy number obtained from aliquots of two independent DNA-libraries made for each sample. “scan”: scanned aliquot, “no scan”: non-scanned control, “EB”: extraction blank, “LB”: library blank. Copy numbers were normalised by the amount of extract included in each library and mean copy numbers were calculated from both libraries. Values and standard deviation are shown on a logarithmic scale.

### Effect of increasing X-ray dose on aDNA quantity

To investigate a possible correlation between an increasing radiation dose and its effect on aDNA, and to define what level of irradiation could be considered harmful for future aDNA analysis, aliquots from different bones were exposed to increasing dose levels, from 0 to 93.2 kGy, by changing exposure time from 0 to 1000 s. The estimated number of total library molecules from each sample was normalised by the amount of extracted material and the corresponding non-scanned aliquot. Mean values were then calculated for each exposure time group. Our results indicate a negative correlation (R^2^ = 0.52) between increasing radiation dose and the number of amplifiable aDNA molecules (Fig. 2).

**Figure 2.**
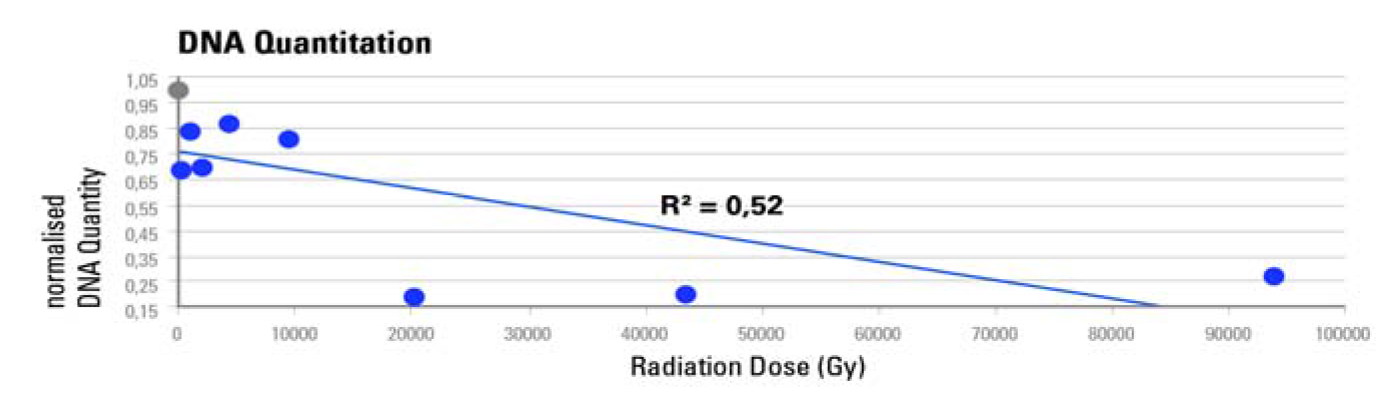
qPCR-based DNA quantitation after irradiation from 0 to 93.72 kGy. Normalised mean aDNA amounts are plotted against cumulative radiation dose received after 0 s (0 Gy), 2.2 s (~206 Gy), 10 s (~937 Gy), 21.5 s (~2.02 kGy), 46.4 s (~4.35 kGy), 100 s (~9.37 kGy), 215.4 s (~20.2 kGy), 464.2 s (~43.5 kGy) and 1000 s (~93.72 kGy). The non-scanned control is represented as a grey circle, the scans as blue circles. Values were normalised by the amount of extracted material and the corresponding non-scanned controls. Mean copy numbers were then obtained per exposure time group consisting of two aliquots from independent DNA-libraries.

### Effect of increasing X-ray dose on aDNA molecule length

In living organisms X-ray radiation is known to induce double strand breaks (DSBs)^22^. Average DNA molecule length is thus assumed to become shorter after radiation exposure. To test if the same effect applies to ancient molecules, we performed mitochondrial capture^23^ and sequenced all enriched libraries from samples that were previously exposed to dose from 0 to 43.5 kGy. No DNA could be sequenced from the 1000 s (93.72 kGy) sample. Sequence data were preprocessed, filtered, and mapped to the mitochondrial reference sequences of the corresponding organisms. Between 4,208,541 and 2,425 fragments mapped to the mitochondrial DNA (mtDNA) reference sequences of the corresponding organism. Mean fragment lengths were determined for mapped (endogenous) and overall (including non-mapped) fragments, and samples were again normalised as described above. The calculated fragment length values were plotted against radiation dose. We observe a strong correlation indicating an almost linear decrease of endogenous (R^2^ = 0.85; Fig. 3a) and overall DNA (R^2^ = 0.66; Fig. 3b) fragment length with increased radiation dose.

**Figure 3.**
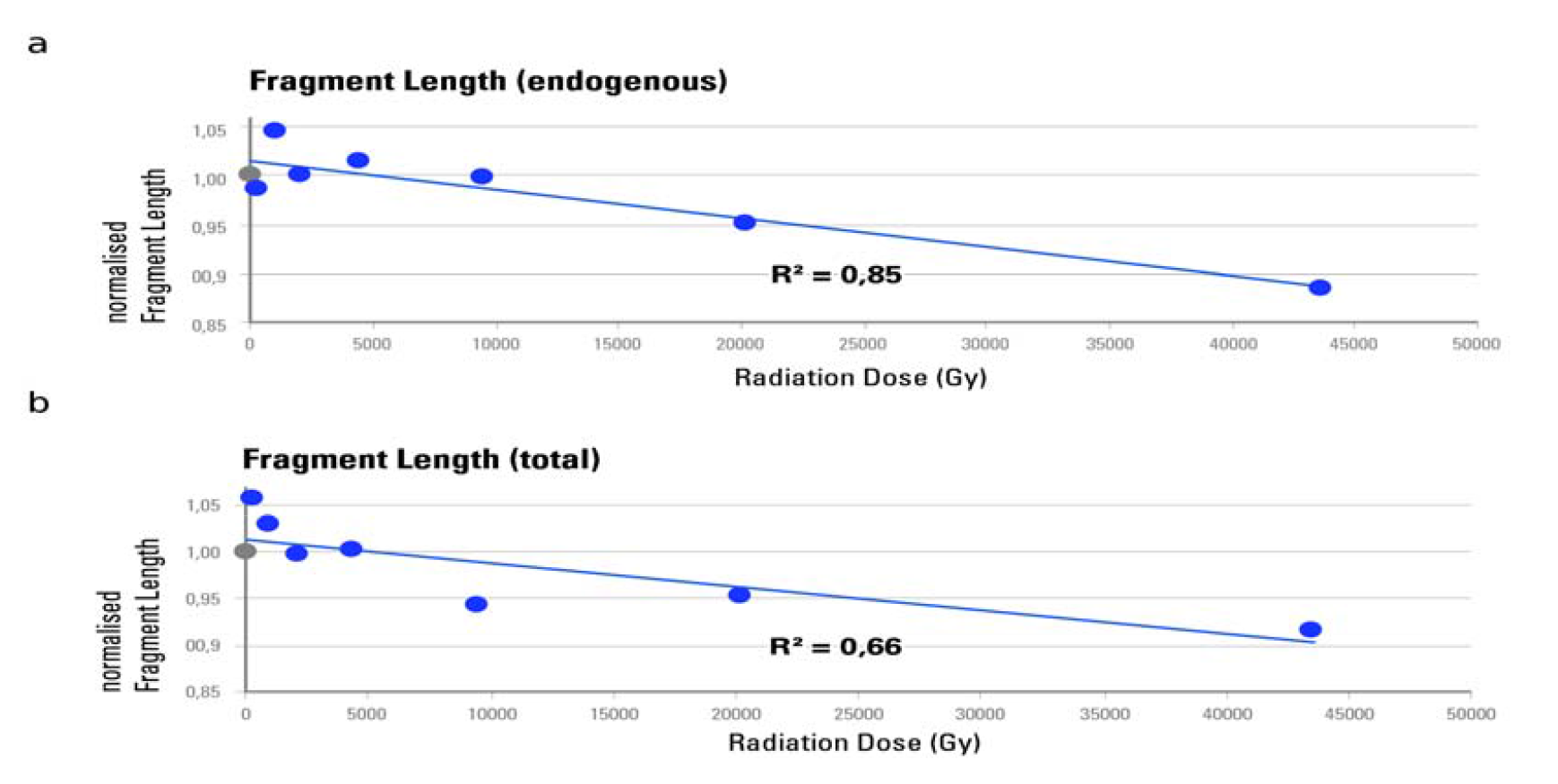
aDNA fragment lengths determined after irradiation from 0 to 43.5 kGy. Normalised mean fragment lengths are plotted against cumulative radiation dose received after 0 s (0 Gy), 2.2 s (~206 Gy), 10 s (~937 Gy), 21.5 s (~2.02 kGy), 46.4 s (~4.35 kGy), 100 s (~9.37 kGy), 215.4 s (~20.2 kGy) and 464.2 s (~43.5 kGy). The non-scanned control is represented as a grey circle, the scans as blue circles. **a.** Mean fragment lengths are calculated for endogenous aDNA fragments only, as determined by mtDNA mapping. **b.** Mean fragment lengths are calculated for total DNA including those fragments that did not map to the corresponding mitochondrial reference sequence. Values were normalised using the means of the corresponding non-scanned control aliquots, and averaged per exposure time group consisting of two aliquots from independent DNA-libraries.

### Increasing X-ray dose and aDNA misincorporation patterns

Since DNA repair mechanisms are absent in dead organisms, chemical modifications of DNA will accumulate. The most common type of such DNA damage is the deamination of cytosines into uracils that causes a nucleotide misincorporation during amplification evident as a C to T substitution most prominently at the 5’ terminus of sequenced aDNA molecules^24^. It has been suggested that the amount of C to T substitutions at the 5’ terminus accumulates over time, and that this can be used to authenticate ancient DNA^25,26^. When DNA molecules get exposed to radiation strong enough to induce strand breaks, the proportion of molecules with terminal substitutions should theoretically decrease, contributing to an associated decrease in the overall C to T substitution frequency (Fig. 4a). A destructive effect of Xray radiation can thus be inferred by reduced terminal substitution frequencies.

To measure the effect of different X-ray doses on nucleotide misincorporation patterns, C to T substitution frequencies were measured for the first position from the 5’ end from the DNA sequence of the exposed aliquots. Data were normalised by the corresponding non-scanned control aliquots and plotted against radiation dose (Fig. 4b). Predictably, substitution frequencies correlate negatively with exposure time, suggesting that a higher X-ray dose lowers the amount of C to T nucleotide misincorporations. This is best explained by DNA strand breaks induced by radiation that introduce new 5’ ends, thus lowering the overall proportion of molecules with terminal C to T substitutions.

**Figure 4.**
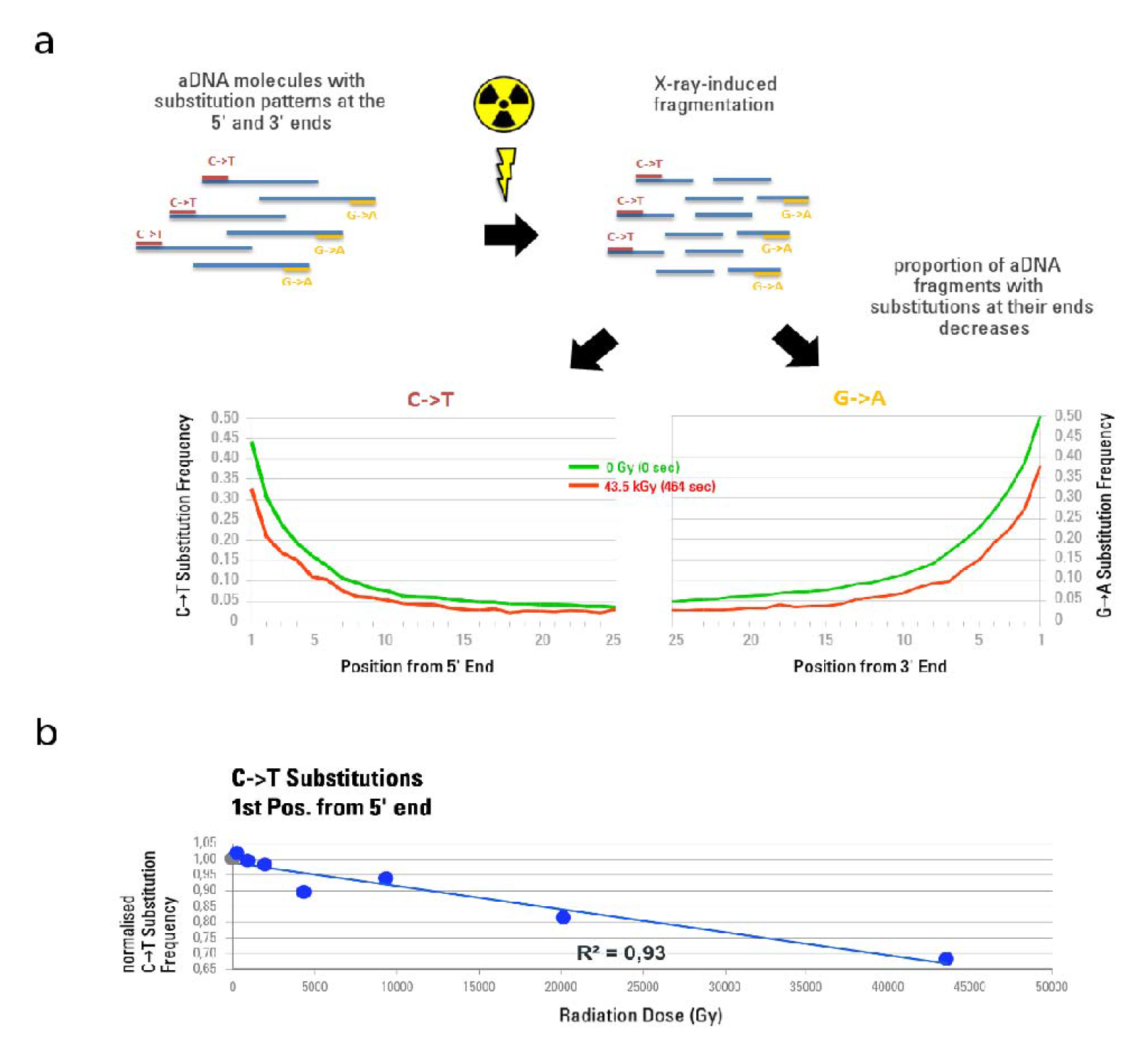
X-ray-induced effects on aDNA nucleotide misincorporation patterns for the first 25 positions from both ends of the molecule. **a.** The misincorporation frequencies of the non-scanned controls are shown as a green line. Lower frequencies (red line) of C to T substitutions at the 5’ ends and G to A substitutions at the 3’ ends can be observed after irradiation at 43.5 kGy, since the fraction of aDNA fragments without misincorporations increases through exposure. **b.** Normalised C to T substitution frequencies of the 1^st^ position from 5’ end of endogenous DNA molecules are plotted against cumulative radiation dose received after 0 s (0 Gy), 2.2 s (~206 Gy), 10 s (~937 Gy), 21.5 s (~2.02 kGy), 46.4 s (~4.35 kGy), 100 s (~9.37 kGy), 215.4 s (~20.2 kGy) and 464.2 s (~43.5 kGy). The non-scanned control is represented as a grey circle, the scans as blue circles. Substitution frequencies were normalised by the corresponding non-scanned controls, and averaged per exposure time group consisting of two aliquots from independent DNA-libraries.

### Effect of synchrotron high quality imaging on aDNA

The results of the first two experiments show no representative effect below 200 Gy, and nearly no effect up to 2 kGy. In order to better assess effects in this dose range, we included three aliquots from cave bear specimens in a series of real μCT scans performed with a classical high quality synchrotron setup (see Material and Methods: Table 2, and Supplementary Information). The total water equivalent surface dose delivered during this scan was 720 Gy, but with an average energy higher than for the previous experiments (127 keV instead of 75 keV). Compared to their non-scanned controls, two out of three libraries showed an increase in the number of amplifiable library molecules after X-ray exposure (Fig. 5a). This may be explained by DSBs introduced by radiation that increase the number of molecules that can be measured via qPCR post ligation of adapters. Even though the same amount of fluorescence signal should theoretically be detected if a molecule was split into multiple fragments, addition of the 132 bp of adapter to each molecule during library preparation will almost double the number of base pairs for a typical aDNA fragment that would be detected by SYBR Green dye intercalation, thus leading to a higher fluorescence signal. One of our samples, Sample 20, however, showed a decrease in aDNA quantity by about 60%. Since this sample derived from a different environmental context (different physicochemical factors) than the other two samples subjected to this test, preservation state of the sample, especially with regard to the level of mineralization and the amount of water exposure, may influence the proportion of aDNA molecules that could be fragmented via the exposure to radiation. An initially higher level of fragmentation for this sample is suggested by its shorter mean read length in the non-scanned aliquot compared to Samples 21 and 22 (Fig. 5b). Further fragmentation due to radiation exposure could have shortened fragments to the point where they would be lost during the purification steps performed during aDNA extraction.

Libraries from all three samples were captured for mtDNA, sequenced, and mapped to the cave bear mitochondrial reference. Since sample 21 remained with only 127 reads after the mapping step, it was not considered further as a threshold of at least 1000 reads was required for further analysis. No consistent trend could be detected for changes in mean fragment length between scanned and non-scanned fractions of samples 20 and 22 (Fig. 5c). In contrast, all samples showed a decrease in mean fragment length as a result of scanning when total reads were considered, applying the same filtering criteria for fragment length and sequence quality (Fig. 5b). We also find that samples 20 and 22 have lowered C to T misincorporation frequencies for the 5’ terminal nucleotide positions after scanning (Fig. 5d).

**Figure 5.**
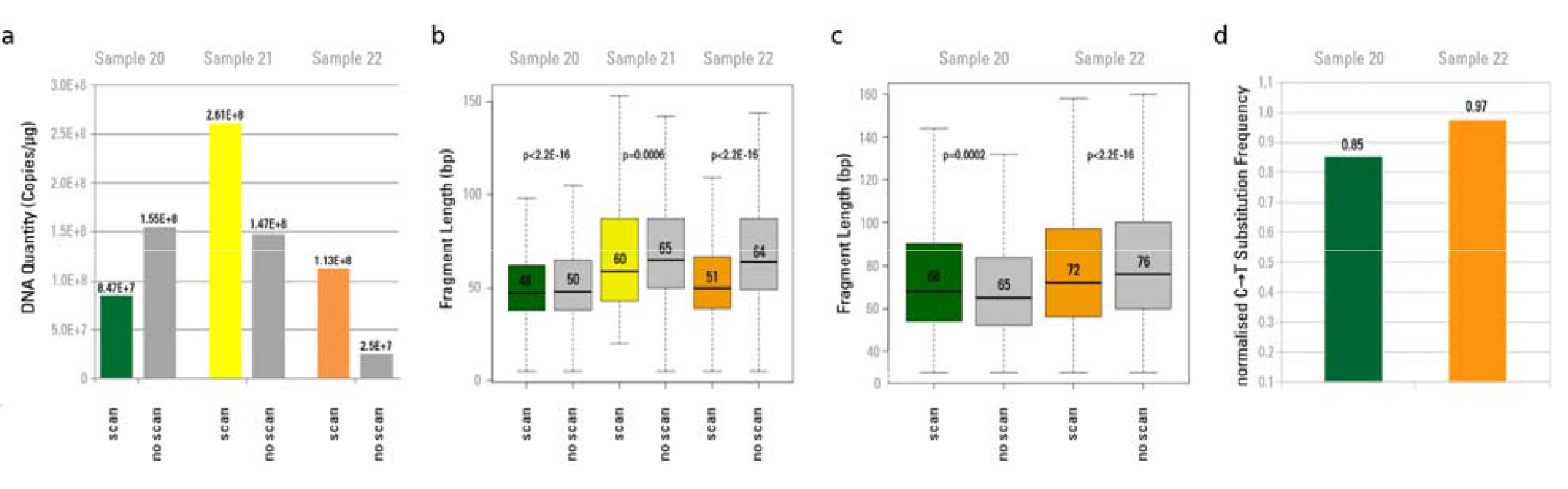
aDNA quantitation, fragment length and misincorporation pattern analyses after high quality imaging synchrotron scan. **a.** aDNA quantitation after exposure to 720 Gy during a real synchrotron μCT scan. “scan”: number of aDNA molecules after scanning the aliquot with 720 Gy, “no scan”: non-scanned control. Values were normalised by the amount of extracted material. **b.** aDNA fragment lengths after exposure to 720 Gy (“scan”) and non-scanned controls (“no scan”). For each aliquot boxplots were generated from fragment lengths of the total DNA content including those fragments that did not map to the target organism’s (here: cave bear) mtDNA. P-values were obtained for each pair of scanned and non-scanned aliquots using Student’s t-test to assess significant differences in mean fragment lengths. **c.** Boxplots were generated only from endogenous (mapped) aDNA molecule fragment lengths. Sample 21 was discarded because of a low number of mapped reads (< 1000). **d.** C to T substitution frequencies of the 1^st^ position from the 5’ end of endogenous aDNA molecules mapped against cave bear mtDNA after X-ray exposure to 720 Gy. The values were normalised by their corresponding non-scanned controls.

### Evaluation of X-ray dose dependent aDNA degradation

Among those samples tested for different exposure times, we cannot observe any effect for doses below 200 Gy. The observed effects for doses between 200 and 2000 Gy are either not detectable or negligible. Above this dose level, we observe significant effects, but they would not preclude successful aDNA analysis for a dose up to 10 kGy. Above this dose level, we observe more severe effects with a rapid decrease in amplifiable DNA, and dramatic effects for doses above 100 kGy that would make aDNA analysis likely impossible (Table 1).

**Table 1.** Summary of effects of X-ray dose on aDNA quantity, molecule length and C to T misincorporation frequencies.

| surface dose (Gy) | aDNA quantity ratio | fragment length ratio endogenous DNA | fragment length ratio total DNA | C/T misincorporation ratio | general effect of X-ray irradiation on aDNA |
| --- | --- | --- | --- | --- | --- |
| 0 | ~1 | ~1 | ~1 | ~1 |  |
|  |  |  |  |  | no detectable effect |
| 200 | ~1 | ~1 | ~1 | ~1 |  |
|  |  |  |  |  | negligible damages |
| 2000 | 0,85 | ~1 | ~1 | 0,97 |  |
|  |  |  |  |  | acceptable damages |
| 10000 | 0,8 | ~1 | 0,98 | 0,93 |  |
|  |  |  |  |  | significant damages |
| 20000 | 0,35 | 0,95 | 0,95 | 0,85 |  |
|  |  |  |  |  | serious damages |
| 45000 | 0,3 | 0,88 | 0,92 | 0,67 |  |
|  |  |  |  |  | strong damages |
| 170000 | 0,1 | N.A. | N.A. | N.A. |  |
|  |  |  |  |  | no usable DNA left |

Coloured lines represent the evaluated risk for the interval between two dose levels. Shown are normalized values corresponding to each applied X-ray surface dose. Normalisation was done by the corresponding non-scanned aliquot. A lower value indicates a more deleterious effect. No effect for dose below 200 Gy could be detected.

### Risk assessment for X-ray imaging and future aDNA analyses

We performed dosimetry of the most relevant configurations used for (μCT on sub-fossils with both conventional and synchrotron sources. In parallel to all the work on aDNA and dosimetry, substantial efforts were also conducted to reduce as much as possible the X-ray dose for synchrotron experiments performed at the ESRF thanks to optimisation of detectors, beam properties and processing protocols. Dosimetry on imaging systems was then performed in three different steps: classical synchrotron experiments, low dose synchrotron experiments and conventional microtomographs.

All measurements and synthesis of results can be found in supplementary information (Supplementary Data 1 and 2). By compiling all the measurements obtained for the conventional microtomographs, we propose a water equivalent surface dose estimator relevant for most of the conventional X-ray imaging systems used to image sub-fossils (Supplementary Data 3).

By combining the results from aDNA analysis and X-ray imaging devices dosimetry, it is possible to evaluate the risk level for future aDNA analyses for both synchrotron scanning and conventional (μCT scanning (Supplementary Data 2).

From our results it is clear that classical synchrotron configurations can indeed reach dangerous levels for aDNA, especially when working at voxel sizes below 10 μm (Fig. 6). Nevertheless, the new systems developed on the ID19-beamline at the ESRF since 2013 put all the configurations used for palaeoanthropology below the detection limit of any defect on aDNA, except the sub-μm resolution setup used for enamel microstructure (Supplementary Information, Note 1, Supplementary Figure 2).

**Figure 6.**
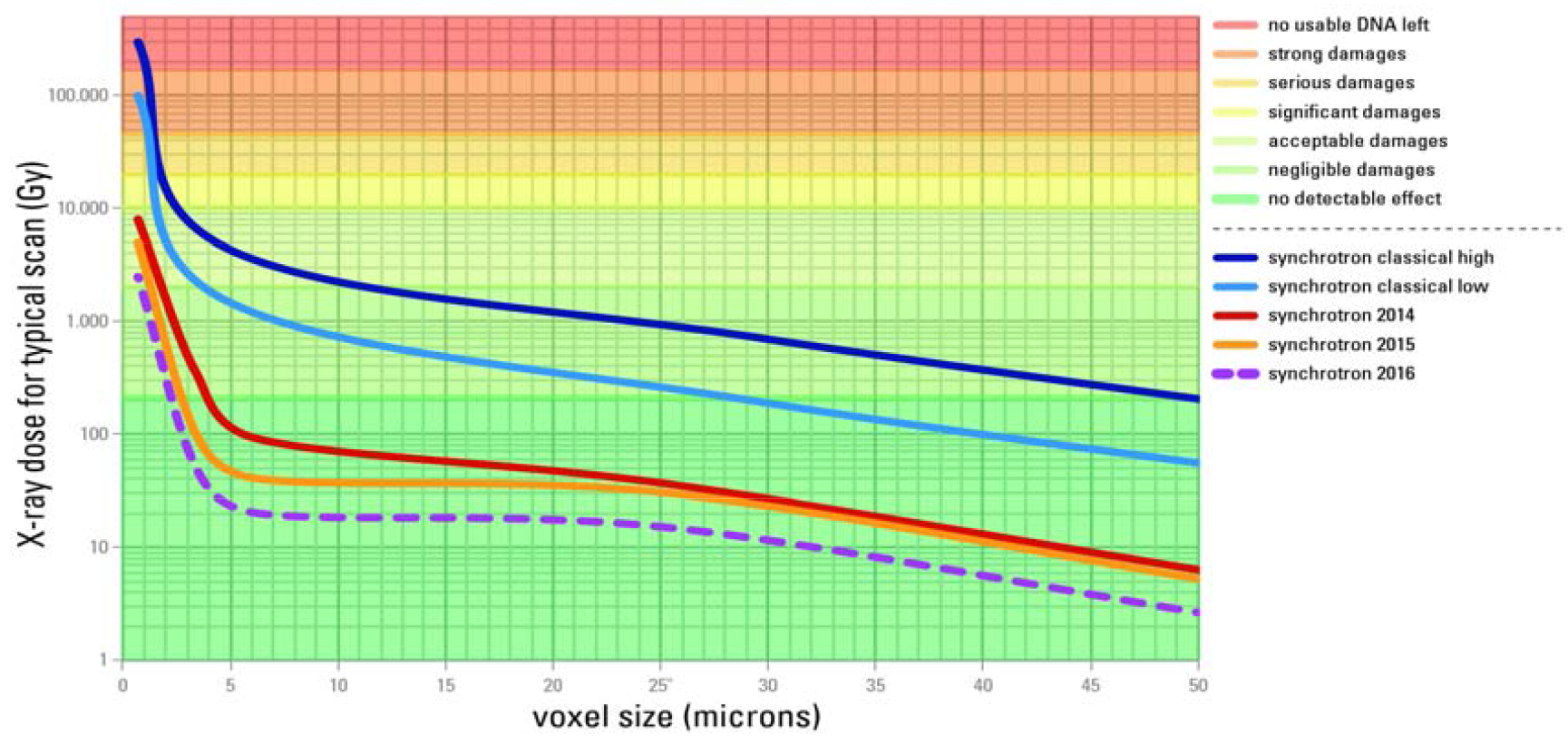
Synchrotron X-ray dose and associated aDNA damages. Typical water surface equivalent X-ray dose per scan and associated damages to aDNA depending on voxel size for synchrotron tomography configurations at the ESRF. Thanks to strong efforts to reduce dose since 2013, the average dose level was reduced by factor 30 in 2014, a factor 62 in 2015, and has reached a factor 125 in early 2016.

It should be noted that these dose values were measured for the ESRF configurations on the ID19-beamline, where much optimisation has been performed during the last 15 years to image fossils. Furthermore, additional optimisation has since been performed to reduce X-ray dose for sub-fossil imaging. These optimisations do not reflect what is available at other synchrotron light sources. Careful dosimetry, setup and spectrum optimisation should therefore be performed on other beamlines before scanning sub-fossil remains that could be potentially subject to aDNA studies.

Dosimetry experiments on the two conventional microtomographs at the MPI-EVA allowed for the assessment of corresponding dose levels for typical scans and to infer the potential risks of aDNA degradation (Fig. 7).

**Figure 7.**
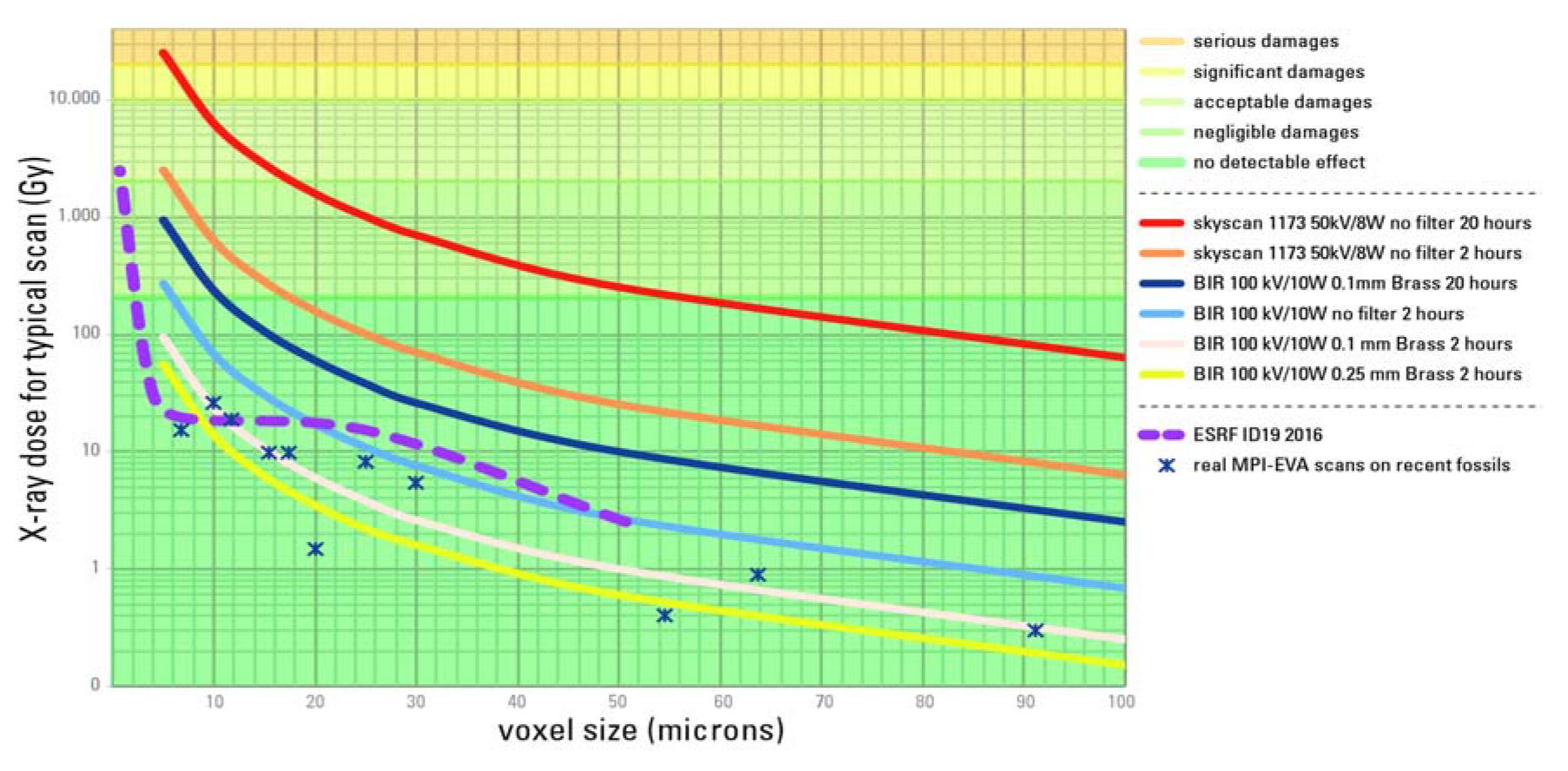
Conventional μCT X-ray dose and associated aDNA damages. Typical water surface equivalent X-ray dose and associated degradation level of aDNA for conventional μCT experiments depending on source parameters, filters, voxel size and scan duration. Plain curves represent typical scanning settings at the MPI-EVA and the ESRF. The dashed curve for ESRF ID19-beamline is calculated based on dose measurements to show the effect of using no filter or thin filter. Only scans without filters, especially repetitions of such scans, can lead to substantial degradation of aDNA. The dose profile of ESRF current configurations is given for comparative purpose.

Scans that are performed on sub-fossils with conventional machines are well below the detection limit of any defects on aDNA. Only very long or repeated scans without any metallic filter with voxel sizes lower than 20μm can endanger aDNA integrity. High surface doses obtained for scans without any filters are due mostly to low energy X-rays. Since the lowest energies of the X-ray spectrum would enhance the beam hardening effect (stronger absorption of the lowest energies in the X-ray spectrum by the sample, leading to higher average energy of the X-rays after the sample than before, resulting into artefacts in the reconstructed data, see Supplementary Information, Note 1, Supplementary Figure 1), a filter of at least 0.1 mm of copper or brass should be used by default for any scan of sub-fossils, thereby providing a higher data quality, while avoiding aDNA-degradation even in case of multiple scans.

A bibliographic survey (Supplementary Table 1) as well as estimation of delivered doses for several specimens scanned at the MPI-EVA before aDNA sampling (Supplementary Information, Supplementary Table 2) shows that in most cases the delivered doses are well below the detection limit of any defect on aDNA, and that samples that were scanned before aDNA sampling do not seem to be different in aDNA-retrieval success rate compared to non-scanned samples.

## Discussion

X-ray-induced damage to aDNA is not yet well understood. Two mechanisms have been suggested. Oxidative damage, either directly caused by ionising radiation or mediated through water radicals, may lead to strand fragmentation or nucleotide modifications including hydantoins, which block DNA polymerases and hence prevent molecules from being amplified^4,5,16,27,28^. Alternatively, X-ray-induced creation of free radicals such as hydroxide may lead to single and/or double strand breaks that may reduce DNA into smaller fragments^14,16,22,27^. In the event of strand fragmentation, an apparent loss of DNA is expected as smaller fragments are assumed to be lost during the purification steps in the DNA extraction. Previous studies have attempted to assess the mechanisms of radiation-induced damage through PCR and electrophoresis-based analyses, and have generated contradictory results in terms of effects on DNA fragmentation^19–21^ (19–21). The samples used for these investigations, modern bones and preserved museum soft tissue such as skin specimens, may not be ideal proxies for evaluating the effect of radiation to aDNA in ancient bones, and although effects of X-radiation on aDNA were recently calculated *in-silico*^18^, the simulations were lacking support from real data obtained from authentic aDNA.

Here we used skeletal remains of late Pleistocene megafauna such as cave bear, giant deer, bison, as well as roe deer to further investigate X-ray-induced effects on authentic aDNA. We used a synchrotron beam for irradiation instead of conventional sources in order to better control all the parameters of the beam, and to be able to cover a large range of dose level without reaching extremely long exposure time. In order to have results relevant for both synchrotron and conventional X-ray sources, we used a polychromatic beam covering an energy range relevant for both kinds of source (Supplementary Information, Note 2, Supplementary Figures 3 - 5). We observed an almost complete loss of amplifiable aDNA in sample aliquots exposed to a high X-ray dose of 170 kGy compared to non-scanned controls. Furthermore, from our second experiment, we observed a decline in aDNA quantity as inferred from qPCR data that correlated with an accumulation in X-ray dose. An almost linear trend was observed in a decrease of mean aDNA fragment length with an increasing radiation dose for endogenous as well as for total DNA molecules. We also found a linear decrease in C to T substitution frequencies at the 5’ end of the DNA fragments with increasing radiation dose. This is best explained by strand fragmentation, where new 5’ ends created by radiation will have a lower frequency of the characteristic damage observed at the ends of ancient DNA fragments. Since this type of nucleotide misincorporation pattern is frequently used to authenticate aDNA molecules^29^, this trend is deserving of special attention, especially with regard to fossils that underwent high irradiation level during their scanning history. The observed damage seems to be proportional to a cumulative X-ray dose, an effect that was previously suggested for modern DNA^30^.

A large amount of DNA defects induced by X-rays in biological conditions are due to the production of extremely reactive free radicals mostly coming from water molecules. Without water, secondary chemical reactions due to free radicals should not take place when scanning dry fossils. Most likely, only direct interactions between X-rays and aDNA molecules (e.g. from direct beam and scattering effects) would have a significant impact. In their analysis of desiccated ancient soft tissues, Paredes *et al.* (2012) did not observe any damaging effects of μCT scanning on DNA^21^, whereas Goetherstrom *et al.* (1995) and Grieshaber *et al.* (2008) purported to have seen X-ray-induced DNA-damage in fresh modern pig bones^19,20^. Based on these data, the amount of water molecules most probably plays a significant role where a higher number of OH-radicals generated in non-desiccated material would ultimately contribute to an increased level of X-ray-induced DNA fragmentation^31^. Strong support for this assumption is given through simulations provided by Wanek and Rühli (2016) who calculated a higher probability for radiation-induced aDNA damage to wet objects based on a higher radical yield, whereas radiation effects on dry objects were non-significant^18^.

In the case of three samples irradiated during a real tomography experiment (Experiment 3), we observed an apparent increase in aDNA molecules after X-ray exposure with a dose of 720 Gy at high energy in two out of three samples. This is best explained by strand breaks that increased the number of DNA molecules in our library, and thus increased the fluorescence signal detected in our qPCR assay. This is further supported by an observed consistent decrease of mean fragment length for total DNA molecules. Despite this decrease in mean length for total DNA, no consistent decline in mean fragment length for endogenous captured cave bear mtDNA could be detected, and the aDNA specific C to T substitutions were only slightly lower in the scanned aliquots. Our application of additional stringent filtering criteria to retain only endogenous molecules, such as mapping to a reference and applying a minimum mapping-quality-filter, may have ultimately reduced the total number of reads subject to analysis, with the shorter or heavily damaged fragments having been removed. Another possibility is that DNA stemming from ancient bone is better protected from ionising radiation than surface contaminants since endogenous DNA will have a higher binding affinity to hydroxyapatite and collagen^16,32^. Hydroxyapatite forms a complex with DNA molecules^33^, and components of the bony matrix might potentially absorb penetrating radiation, leading to a partial shielding. Higher fractions of DNA have been recovered from hydroxyapatite than from collagen^34,35^, emphasizing the importance of the mineral fraction of bone for the preservation of aDNA. Finally, endogenous aDNA is more protected from water than contaminating DNA, and therefore less affected by secondary damages due to free water-radicals.

The results obtained for this configuration during real high quality classical synchrotron scan indicate that for surface doses below 2000 Gy, the effects on aDNA would not lead to substantial degradation and would not compromise palaeogenetic investigations.

## Conclusion

In summary, our results confirm that X-ray irradiation of sub-fossils can have a detrimental effect on aDNA integrity when the total water equivalent surface dose exceeds 200 Gy; the degradation increase being roughly linear with the dose accumulation. While the effect is very limited up to 2 kGy, the degradation can reach dramatic levels for doses exceeding 100 kGy. Based on the results presented above, we have defined the detection limit at 200 Gy due to the limited number of samples irradiated in the 200-720 Gy dose range. Nevertheless additional experiments would be necessary to define this limit more precisely. In order to ensure safe scanning, we therefore suggest the following guidelines for the scanning of sub-fossil remains:

### Recommendation for conventional μCT of sub-fossils

- Never perform scans without a metallic filter. Always use filters to remove lowest energies of the spectrum. We recommend a systematic use of at least 0.1 mm of copper or brass filter for scanning of sub-fossils. Such a filter will ensure safe scanning while increasing data quality by reducing the beam hardening.

### Recommendations for synchrotron scanning of sub-fossils

- As synchrotron scanning can clearly lead to aDNA degradation, careful dosimetry has to be performed before any real experiment on a synchrotron beamline involving scanning of sub-fossils. Only high optimisation effort can put synchrotron setup in the safe dose region.
- Perform dose estimation/measurements before doing the actual scan. Take into account overlap of scans.
- Phase contrast being up to 1000 times more sensitive than absorption one, always use it to optimise results while keeping the dose as low as possible.
- Always perform a precise collimation of the beam (beam size fitting the field of view) to ensure that as low dose as possible is delivered to non-imaged areas.
- For sub-μm resolution scans, orient the samples carefully to minimize the amount of material crossing the beam path.

### Recommendations for sub-fossil scanning in general

- Never scan a wet specimen.
- Do not perform scans at higher resolution than necessary, as the dose increases roughly at the square power of the increase of resolution.
- Dose is cumulative: do not perform multiple scans when similar data already exist. Data sharing (under responsibility of the curators) and public repository databases play an important role on this.
- Record precise information about all scanning parameters and scanning geometry and provide them to curators in order to track the whole scanning history for a given specimen.
- Before any new scanning of a specimen, take into account its scanning history to estimate the total dose accumulation (especially in the case of synchrotron experiments).

These simple recommendations can ensure that both conventional and synchrotron X-ray scanning of sub-fossils will not hinder future aDNA investigations and still allow studying the morphological or microstructural information by X-ray imaging before destructive sampling.

## Material and Methods

### Samples

Since sub-fossilised human remains were too rare to be used for this study, we collected animal samples considered old enough to be representative of ancient human bones, such as Neanderthals. In total, a maximum of 38 samples from ancient faunal remains could be obtained. The selected species represent cave bear (*Ursus spelaeus*), giant deer (*Megaloceros giganteus*), bison (*Bison priscus*) and roe deer (*Capreolus capreolus*) which were obtained from the Swabian Jura, the French Jura and the Ukraine. The samples were UV-irradiated overnight to remove surface contamination, crushed into 0.5-2.0 mm^2^ fragments and subdivided into 200 mg aliquots. To ensure non-biased results, a double-blind approach was used: no information about aliquot origin and preservation state was given to the persons in charge of the synchrotron scanning. As well, no information concerning the scanning settings and the organisation of scanned aliquots and control aliquots was available for the persons in charge of the aDNA extraction and quantification.

### X-ray exposure experiments and dosimetry

#### 1. Synchrotron (ESRF)

All X-ray exposure experiments were conducted on the BM05 beamline at the European Synchrotron Radiation Facility (ESRF) in Grenoble, France. The dose rate for the various setups was measured using a PTW Unidos E (T10021) dosimeter equipped with a TM31010 semiflex ionisation chamber. Measurements were conducted as surface water equivalent dose, the ionisation chamber sensitive part being completely covered by the X-ray beam. The total integrated dose for a complete scan was then calculated as the total exposure time multiplied by the dose rate (Table 2).

**Table 2:**
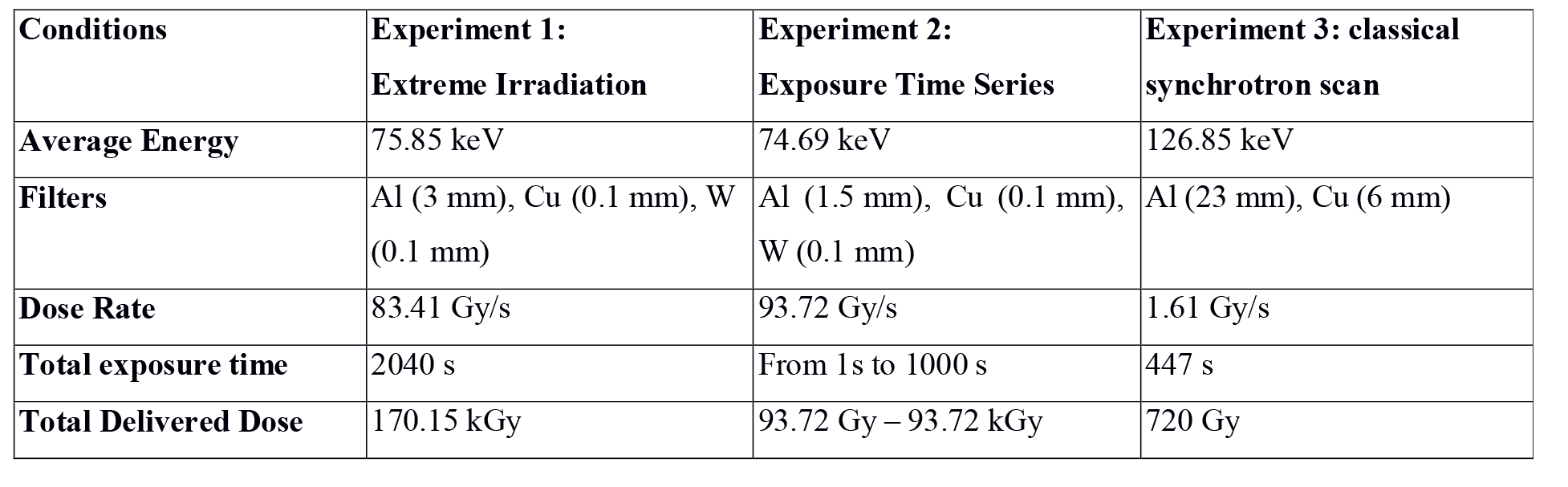
Overview of the used conditions during the three conducted experiments.

| Conditions | Experiment 1:<br>Extreme Irradiation | Experiment 2:<br>Exposure Time Series | Experiment 3: classical<br>synchrotron scan |
| --- | --- | --- | --- |
| Average Energy | 75.85 keV | 74.69 keV | 126.85 keV |
| Filters | Al (3 mm), Cu (0.1 mm), W (0.1 mm) | Al (1.5 mm), Cu (0.1 mm), W (0.1 mm) | Al (23 mm), Cu (6 mm) |
| Dose Rate | 83.41 Gy/s | 93.72 Gy/s | 1.61 Gy/s |
| Total exposure time | 2040 s | From 1s to 1000 s | 447 s |
| Total Delivered Dose | 170.15 kGy | 93.72 Gy – 93.72 kGy | 720 Gy |

In a first experiment, aliquots from 11 different samples were irradiated every 45 degrees over 360 degrees using a polychromatic beam (50 × 5 mm^2^) with the following settings: BM05 bending magnet white beam filtered with 3 mm of aluminium, 0.1 mm of copper and 0.1 mm of tungsten leading to an average energy of 75.85 keV (Supplementary Information, Note 2, Supplementary Figure 3); 200 mA in the storage ring, sample at 50 m from the source; 17 min/scan; each part of sample scanned twice. The measured dose rate was 83.41 Gy/s, yielding a total delivered dose at every position in the samples of 170.15 kGy.

In a second experiment, aliquots were scanned using a polychromatic beam (45 × 3.1 mm^2^) with the same settings as described above (except 1.5 mm of Al instead of 3 mm, Supplementary Information, Note 2, Supplementary Figure 4), but at different exposure times (0 s, 1 s, 2.2 s, 4.6 s, 10 s, 21.5 s, 46.4 s, 100 s, 215.4 s, 464.2 s, and 1000 s) and a measured dose rate of 93.72 Gy/s which yielded a maximum delivered X-ray dose of 93.72 kGy. For each different time of exposure, the delivered dose is as follows: 0 Gy, 93.72 Gy, 206 Gy, 431.11 Gy, 937 Gy, 2.02 kGy, 4.35 kGy, 9.37 kGy, 20.19 kGy, 43.5 kGy and 93.7 kGy. A fast shutter was used to ensure that the total exposure time was equally distributed over 360 degrees every 45 degrees.

In a third experiment, aliquots were placed in a plastic tube together with cave bear (*Ursus spelaeus*) mandibles that underwent high quality scanning for other research purposes. Each sample was scanned twice (50% overlap between consecutive vertical scans). Scanning parameters involved a polychromatic beam at an average energy of 127 keV, obtained by filtering the bending magnet beam by 23 mm of aluminium and 6 mm of copper (Supplementary Information, Note 2, Supplementary Figure 5). We used a propagation distance for phase contrast of 2.4 m, and a pixel size of 29.88 μm. The total exposure time for each part of the samples was 447 s, with a measured dose rate of 1.61 Gy/s (dosimeter in the plastic tube), leading to a total delivered X-ray dose of 720 Gy.

#### 2. Conventional μCT scanners (MPI-EVA)

The dosimetry characterisation for conventional μCT-scanners was performed at the Max Planck Institute for Evolutionary Anthropology (MPI-EVA, Leipzig, Germany) on the two portable industrial μCT-scanners: a BIR ACTIS 225/300 and a Skyscan 1173. The dose rate was measured using the same model of dosimeter as for the synchrotron measurements (PTW: UNIDOS Webline T10021, Ionisation chamber TM31010), with the same calibration protocol using a ^60^Co radioactive source. For both scanners, the dose rate was measured in different conditions of irradiation, involving variation in power, voltage, absence or presence of metallic filter of various thickness and material. Ranges of parameters explained below include conditions where the dose rate was not measurable in some cases, but this clearly appears in Supplementary Data 1.

For the BIR, three sessions of measuring were run. Firstly, the dosimeter was fixed on the sample stage at 250 mm from the X-ray source, initially with no metallic filter and then with various filters (0.25 mm Brass, 0.5 mm Brass, 1 mm Brass, 1 mm Al), the operators were then able to measure the dose rate for power ranging from 5 to 150 W and voltage from 50 to 200 kV. Secondly, for each filtering condition mentioned above, the dose rate was measured at 50 kV, 130 kV and 200 kV and placing the dosimeter at an increasing distance from the dosimeter from 20 mm to 500 mm. Last, the influence of distance to beam axis was tested at 130 kV, 50 W, 250 mm from the X-ray source, and with 0.5 mm of Brass. This involved combinations of variations in altitude (z ranging from 0 to 100 mm) and in translation (from 0 to 100 mm). It has to be noted that 130 kV is the most commonly used voltage when scanning at MPI-EVA.

For the Skyscan 1173, the dosimeter was set at the centre of the rotation stage, first at 125 mm from the X-ray source (⇔ 17.2 μm, commonly used distances for scans at MPI-EVA), and measures were taken twice during irradiations without filter, with 1 mm Al, 0.25 mm Brass, with a power ranging from 2W to 8W, and a voltage from 50 kV to 130 kV. Second, with the same range of power, the sample stage was moved from 50 mm to 250 mm to the X-ray source without filter (at 50kV and 80 kV), 1mm Al and 100 kV, and with 0.25mm Brass and 130 kV. Last, the dose rate was measured at 130 kV, with 0.25 mm Brass, at 125 mm from the source, with a power 2-8 W, and with a translation in “z” from 0 to 60 mm.

#### aDNA Extraction and Preparation of Sequencing Libraries (University of Tübingen)

Prior to extraction samples were UV-irradiated overnight. DNA extraction was carried out using 50 mg as starting material based on a guanidinium-silica based extraction method^36^. For each sample a DNA library was prepared according to published protocols^37,38^ using 20 μl of extract. Sample-specific indexes were added to both library adapters to allow differentiation between individual samples after pooling and multiplex sequencing^38^. Indexed libraries were amplified in 100 μl reactions in a variable number of 9 to 14 cycles to reach the amplification plateau, followed by purification over *Qiagen MinElute* spin columns (*Quiagen*, Hilden, Germany).

#### Quantitation of aDNA Amount

qPCR quantification of the libraries was carried out on a Roche LightCycler 480 using the DyNAmo HS SYBR Green qPCR kit by Thermo Scientific. qPCR primers and standards are described in Meyer and Kircher 2010^37^. All DNA quantities were numbered as a number of copies - i.e. a number of fragments with an adapter on each end - per μl of library. As there is no amplification during library preparation, variation in the number of copies between libraries is assumed to reflect the variation in the DNA quantity between extracts. The number of copies for each library was normalised by the amount of extracted material in order to prevent a bias due to sampling. Results were analysed using the *Roche* Lightcycler’s 480 integrated software, Microsoft Excel and the R statistical software^39^. The obtained values were normalised by division through the corresponding value of the non-scanned control aliquot. Except for the last experiment, mean values were calculated per scan group consisting of two aliquots from independent sequencing libraries. See Supplementary Data 4 for normalised DNA quantitations in Fig. 1, Fig. 2, Fig. 3 and Fig. 5.

#### Enrichment of aDNA

Target enrichment of mtDNA was performed by capture of the pooled libraries using bait generated from roe deer (*Capreolus capreolus*), brown bear (*Ursus arctos*) and bison (*Bison bison*) mtDNA^23^. The bait was generated by use of three primer sets (Table 3) designed with the Primer3Plus software package^40^. All extractions and pre-amplification steps of the library preparation were performed in clean room facilities and negative controls were included for each reaction. Globally, all criteria established for ancient DNA studies authenticity were respected.

**Table 3.**
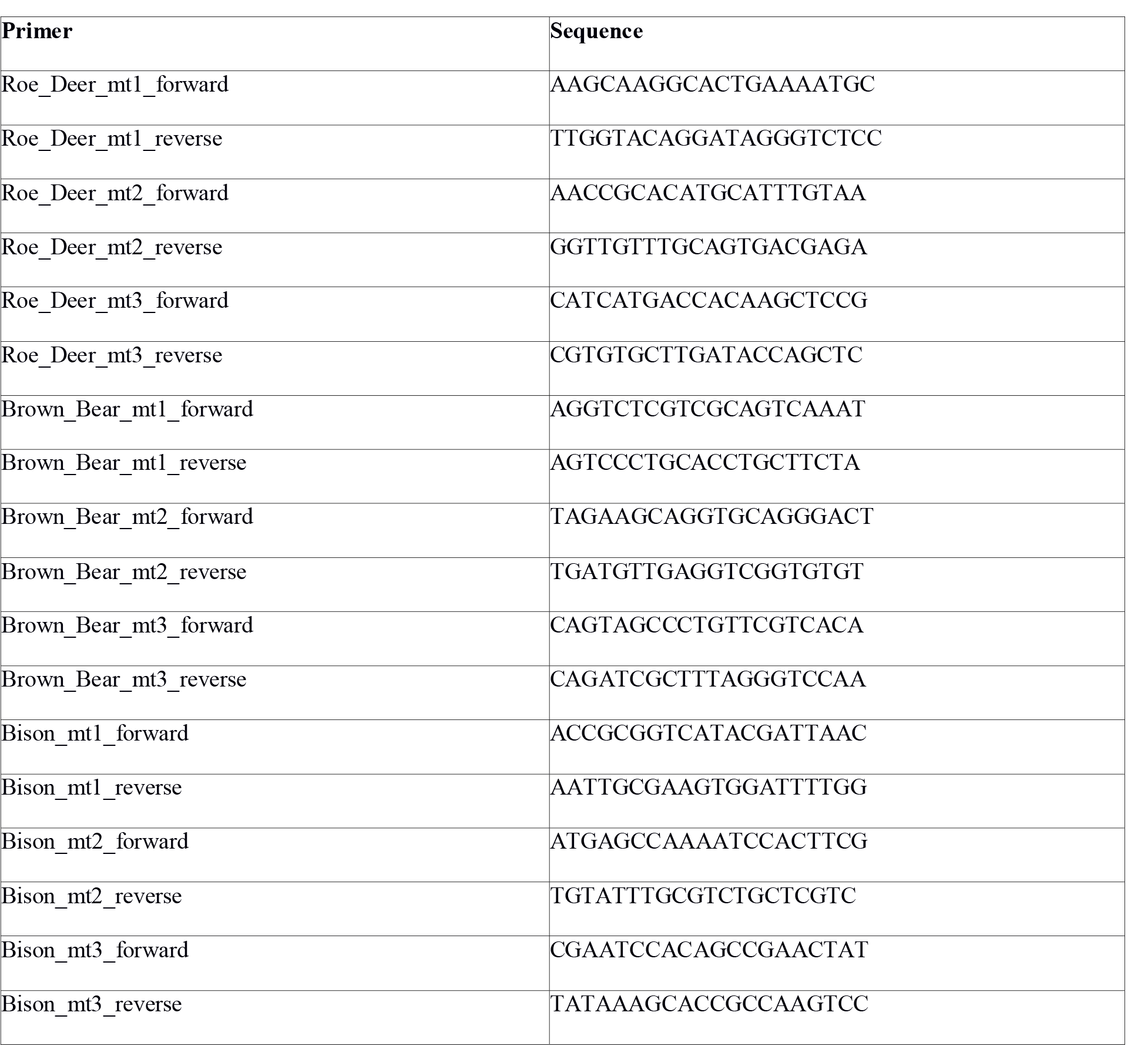
Forward and reverse primer pairs used to generate bait for targeted mtDNA enrichment.

#### Sequencing and Preprocessing of Raw Sequences

Indexed library pools were sequenced on the Illumina Hiseq 2500 platform with 2*100+7+7 cycles^38^. Demultiplexing was performed by sorting all the sequences corresponding to their p7 and p5 index combinations. Forward and reverse reads were merged into single sequences if they overlapped by at least 11 bp^41^. Unmerged reads were discarded and merged reads were filtered for a length of at least 30bp.

#### Mapping

The preprocessed sequences were mapped to one of the corresponding complete mitochondrial genome reference sequences: *Capreolus capreolus* (NC_020684.1), *Bison bison* (NC_012346.1) or *Ursus spelaeus* (FM177760.1). Mapping was performed using BWA^42^ with seeding turned off and a reduced mapping stringency parameter “-n 5” to account for up to five mismatches in ancient DNA reads^43^. Duplicate removal was performed on those reads that showed identical start and end coordinates only. The produced BAM files were filtered for sequences with a mapping quality of at least 20. We set a threshold of at least 1000 mapped reads for an aliquot to be considered in further analyses.

#### Fragment Lensth Distribution Analysis

Read lengths were obtained from quality filtered BAM files using SAMtools^44^. Read length means and boxplots were obtained using the R statistical software^39^. Significance testing was assessed using the unpaired Student’s t-Test in R. Normalisation of the data was achieved by dividing the obtained mean values by the mean fragment length of the corresponding non-scanned control aliquots. Scatter plots for regression analysis were generated in Microsoft Excel. See Supplementary Data 4 and 5 for normalised fragment length values in Fig. 3 and Fig. 5.

#### Analysis of aDNA C to T Substitution Frequencies

C to T misincorporation frequencies typical of aDNA were obtained using mapDamage 2.0^45^. Data was normalised by division through the corresponding value of the non-scanned control for each position from 5’ end. Scatter plots and bar plots were generated in Microsoft Excel. For the exposure time experiment, mean values were calculated per scan group consisting of two aliquots from independent sequencing libraries. See Supplementary Data 4 for the substitution frequencies in Fig. 4 and Fig. 5 d.

## ACKNOWLEDGEMENTS

We thank the ESRF BM05 beamline to have provided access to beamtime, as well as Thierry Brochard for his help with the dosimetry experiments. The computational work was performed on the computational resource bwGRiD Cluster Tübingen funded by the Ministry of Science, Research and the Arts Baden-Württemberg and the Universities of the State of Baden-Württemberg, Germany, within the framework program bwHPC. We want to thank Marek Dynowski for providing access to the computational resource bwGRiD Cluster Tübingen and his technical support. This research was partially funded by the European Research Council Starting Grant APGREID (to A.I., A.H., K.B. and J.K.). We want to thank James Fellows Yates for proofreading the manuscript and improving the language. We thank Cosimo Posth and Maria Spyrou for their helpful comments. We also wish to acknowledge Michel Toussaint for providing access to the Engis 2 Neandertal material. We are grateful to Svante Pääbo for his help and support at an earlier stage of this project.

### Author Contributions

J.K., P.T., M.B., A.I. and A.L.C conceived and designed the general research. A.I., M.B. and J.K. further designed the genetic analyses while P.T. and A.L.C designed and conducted the irradiation-experiments at the ESRF. P.T., A.L.C. and H.T. designed the conventional scanner dosimetry measurements that were performed by A.L.C. and H.T. at the MPI-EVA. M.B., F.L. and V.J.S. performed mtDNA-extraction and the preparation of sequencing libraries at the University of Tübingen. N.J.C., S.C.M., A.B., G.P., D.G.D., M-A.J. and O.K. provided samples. B.V., K.H., P.T. and J-J.H. facilitated the research and provided access to infrastructure. A.I. and A.H. performed bioinformatic analyses at the MPI-SHH. A.I., P.T. and J.K. wrote the manuscript and A.L.C. and K.B. contributed to it. All authors reviewed the manuscript.

### Competing financial interests

The authors declare no competing financial interests.

